# Cannabinoid signaling modulation through JZL184 restores key phenotypes of a mouse model for Williams-Beuren syndrome

**DOI:** 10.1101/2021.08.16.456474

**Authors:** Alba Navarro-Romero, Lorena Galera-López, Paula Ortiz-Romero, Alberto Llorente-Ovejero, Lucía de los Reyes-Ramírez, Aleksandra Mas-Stachurska, Marina Reixachs-Solé, Antoni Pastor, Rafael de la Torre, Rafael Maldonado, Begoña Benito, Eduardo Eyras, Rafael Rodríguez-Puertas, Victoria Campuzano, Andrés Ozaita

## Abstract

Williams-Beuren syndrome (WBS) is a rare genetic multisystemic disorder characterized by mild to moderate intellectual disability and hypersocial phenotype, while the most life-threatening features are cardiovascular abnormalities. Nowadays, there are no available treatments to ameliorate the main traits of WBS. The endocannabinoid system (ECS), given its relevance for both cognitive and cardiovascular function, could be a potential druggable target in this syndrome. We analyzed the components of the ECS in the complete deletion (CD) mouse model of WBS and assessed the impact of its pharmacological modulation in key phenotypes relevant for WBS. CD mice showed the characteristic hypersociable phenotype with no preference for social novelty and poor object-recognition performance. Brain cannabinoid type-1 receptor (CB1R) in CD male mice showed alterations in density and coupling with no detectable change in main endocannabinoids. Endocannabinoid signaling modulation with sub-chronic (10 d) JZL184, a selective inhibitor of monoacylglycerol lipase (MAGL), specifically normalized the social and cognitive phenotype of CD mice. Notably, JZL184 treatment improved cardiac function and restored gene expression patterns in cardiac tissue. These results reveal the modulation of the ECS as a promising novel therapeutic approach to improve key phenotypic alterations in WBS.

## Introduction

Williams-Beuren syndrome (WBS) is a genetic neurodevelopmental disorder caused by a hemizygous deletion of a region containing 26 to 28 genes at chromosomal band 7q11.23. The estimated prevalence of this disorder is 1 in 7,500 individuals (1). Subjects present manifestations affecting mainly the central nervous system and the cardiovascular system (2). WBS subjects show mild-to-moderate intellectual disability with an intelligence quotient (IQ) score from 40 to 90 (3, 4) affecting their quality of life, where independent living is infrequent (5). One of the most prominent features of the cognitive profile of WBS is an hypersociable phenotype characterized by uninhibited social interactions and a reduced response to social threat (6, 7). This phenotype is opposite to the archetypic social phenotype of autism spectrum disorders characterized by lack of social interest and deficits in social communication (8). Notably, the congenital cardiovascular phenotype in WBS is the major source of morbidity and mortality (9), characterized by elastin arteriopathy, supravalvular aortic stenosis, peripheral pulmonary stenosis and hypertension, which require in many cases surgical correction (10). While strategies such as behavioral intervention can improve WBS cognitive skills to some extent, or certainly invasive surgical procedures are available, their success is limited and WBS is largely without treatment (11).

Several mouse models have been developed mimicking the genetic alterations observed in WBS subjects (12). Among them, the complete deletion (CD) mouse model resembles the most common hemizygous deletion found in WBS patients and displays several WBS phenotypic traits (13). Indeed, this model shows significant alterations in social behavior with enhanced sociability (13), cognitive deficits (14) and a mild cardiovascular phenotype with cardiac hypertrophy, borderline hypertension and mildly increased arterial wall thickness (13) among others.

The endocannabinoid system (ECS) is an homeostatic neuromodulator system involved in many physiological functions and behavioral responses including sociability (15) and cognition (16–18). It is composed by cannabinoid receptors, including cannabinoid type-1 and cannabinoid type-2 receptors (CB1R and CB2R, respectively), their endogenous ligands or endocannabinoids (mainly, N-arachidonoylethanolamine, AEA, and 2-arachidonoylglycerol, 2-AG), and the enzymes involved in the synthesis and inactivation of these ligands. Interestingly, alterations of this neuromodulatory system have been described in several mouse models for autism spectrum disorders with altered sociability (19). In addition, the pharmacological modulation of the ECS, whether targeting the cannabinoid receptors, or the enzymes involved in the degradation of the endocannabinoids, restores social abnormalities in some of these models (19, 20).

In this study, we investigated the brain components of the ECS in the WBS CD model to find significant alterations in CB1R expression in particular brain regions. Additionally, we reveal that sub-chronic administration of the monoacylglycerol lipase (MAGL) inhibitor JZL184 normalized relevant behavioral phenotypes in CD mice including social behavior and memory alterations. Interestingly, this treatment also partially restored cardiac transcriptional alterations found in the model. Altogether, our results indicate that the modification of the endocannabinoid signaling could be a novel therapeutic strategy worth evaluating in the context of WBS.

## Results

### CD mice exhibit social and cognitive alterations

We first analyzed social behavior in CD mice and their WT littermates using the Vsocial-maze. (21) (Fig. 1a). This test allows to assess exploration, sociability and preference for social novelty in the same mouse. Exploratory behavior was analyzed in the empty Vsocial-maze during the habituation phase by accounting the time mice explored both empty compartments (E1 and E2) at the end of the corridors. No changes were observed between genotypes (Fig. 1b, left). Then, during the sociability phase, both WT and CD mice displayed a significant preference for exploring a compartment with an unfamiliar juvenile mouse (stranger 1, S1) rather than an empty compartment. Notably, CD mice spent significantly more time than WT mice exploring stranger 1 (Fig. 1b, middle). Finally, during the preference for social novelty phase, CD mice explored similarly stranger 1 and a novel unfamiliar juvenile mouse (stranger 2, S2) in contrast to WT animals that showed a significant preference for the novel stranger (Fig. 1b, right). These data indicated that CD mice presented an hypersociable phenotype and a lack of preference for social novelty.

**Figure 1.**
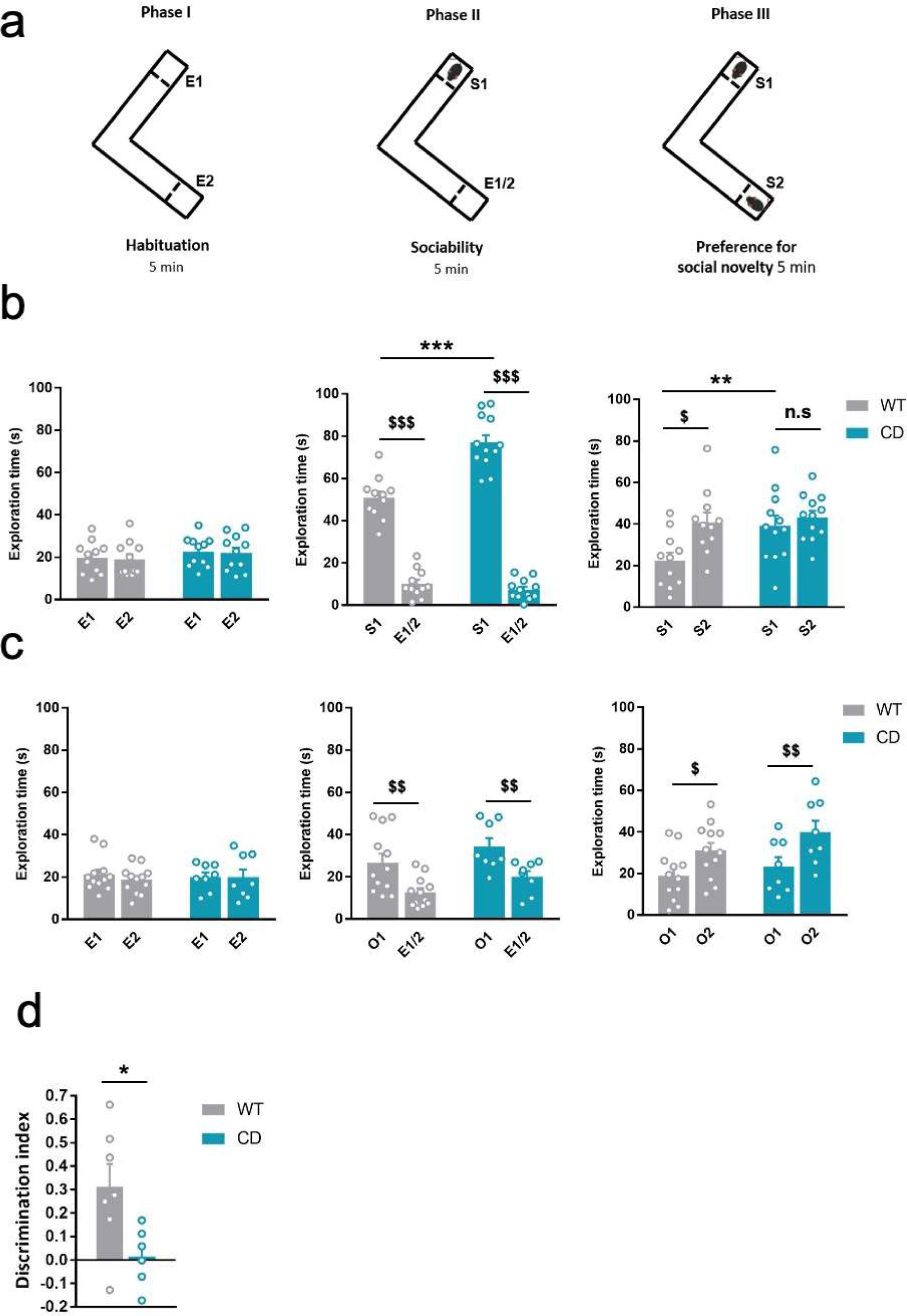
CD mice show an hypersociable phenotype, no preference for social novelty and cognitive alterations. (a) Schematic cartoon of the sociability and preference for social novelty procedure. (b) Time spent exploring either empty compartments (E) or stranger mice (S) during the three phases of the Vsocial-maze (WT, n=11; CD, n=11). (c) Time spent exploring either empty compartments (E) or objects (O) (WT, n=12; CD, n=12). Statistical significance was calculated by repeated measures ANOVA comparison. $ p<0.05; $$ p<0.01; $$$ p<0.001 (compartment effect); ** p<0.01; *** p<0.001 (genotype effect). (d) Discrimination index of WT and CD mice (WT, n=7; CD, n=6). Statistical significance was calculated by Student’s t-test. * p< 0.05 (genotype effect). Data are expressed as mean ± S.E.M.

To confirm that both alterations in social behavior were dependent on social stimuli, we repeated the same procedure using unfamiliar objects instead of unfamiliar mice (see setting in Supplementary Fig. 1). Both WT and CD mice displayed a preference for the compartment with an object (object 1, O1) over the empty compartment. In contrast to social behavior, WT and CD mice spent similar time exploring object 1 (Fig. 1c, middle). When a new object (object 2, O2) was introduced, both WT and CD mice spent more time exploring the new object than the object that had been previously explored (Fig. 1c, right). On the one hand, these results reveal the strong motivation of CD mice for social interaction compared to WT mice. On the other hand, they show both genotypes display similar motivation to explore object novelty. Under our experimental conditions, using the V-maze, CD mice also displayed an impairment in short-term memory in novel-object recognition test (NORT) (Fig 1d).

### CD mice show alterations in density and signaling of CB1R

We determined the levels of endocannabinoids AEA, 2-AG and related N-acylethanolamines (N- docosatetraenoylethanolamine, DEA and N-docosahexaenoylethanolamine, DHEA) and 2- monoacylglycerols (2-linoleoylglycerol, 2-LG and 2-oleoylglycerol, 2-OG) in whole brain homogenates. No significant changes were revealed in CD mice in comparison to WT animals (Table 1). Then, we analyzed cannabinoid receptor brain density by [^3^H]CP55,940 radioligand binding assay in brain tissue sections. Quantitative densitometry revealed an increased density of cannabinoid receptors in the basolateral and central amygdala, whereas a decreased density was observed in the polymorphic and granular layers of dentate gyrus (Fig. 2, a and b) (Supplementary Table 1). To determine the specific subtype of the cannabinoid receptor studied, rimonabant and SR144528, selective antagonists for CB1R and CB2R, respectively, were used. Rimonabant, but not SR144528, blocked [^3^H]CP55,940 radioligand binding in brain slices (Supplementary Fig. 2a) indicating that the observed changes occur in CB1R distribution.

**Figure 2.**
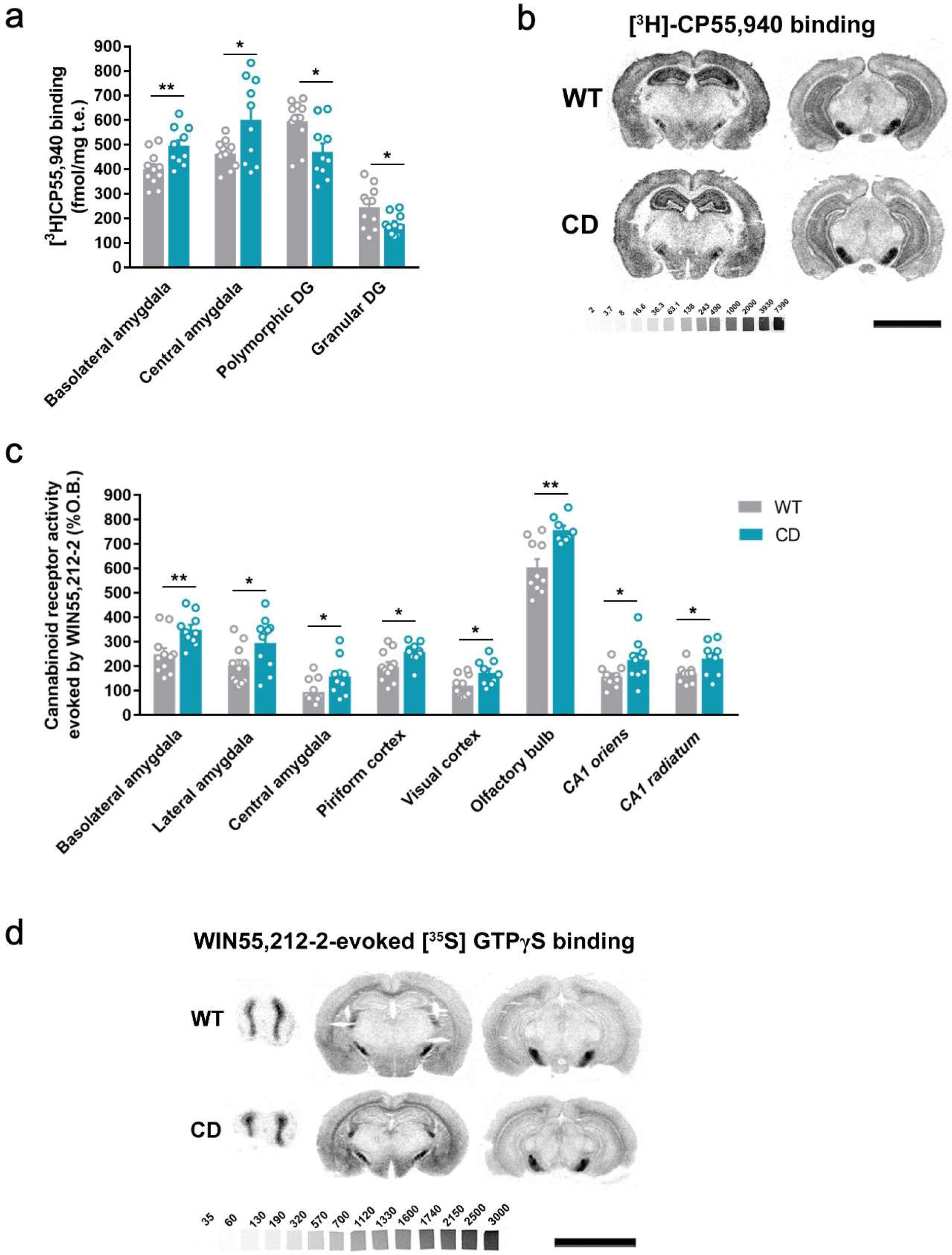
CD mice show alterations in CB1R density and signaling. (a) [^3^H]CP55,940 binding of brain regions with significant changes in CD mice in comparison to WT littermates (WT, n=12; CD, n=10). (b) Representative images of [^3^H]CP55,940 binding autoradiography (c) [^35^S]GTPγS binding evoked by WIN55,212-2 (10 µM) of brain regions with significant changes in CD mice in comparison to WT littermates (WT, n=12; CD, n=10) expressed as percentage of stimulation over the basal binding. (d) Representative images of WIN55,212-2-evoked [^35^S]GTPγS binding. [^14^C]- microscales used as standards in Ci/g t.e. Scale bar = 5 mm. Statistical significance was calculated by Student’s t-test. * p<0.05; ** p<0.01; (genotype effect). Data are expressed as mean ± S.E.M.

**Table 1.** Relative levels of endocannabinoids and related compounds in whole brain homogenates of CD and WT. (WT, n=10; CD, n=11). Statistical significance was calculated by Student’s t-test. Data is expressed as mean ± S.E.M.

|  | Whole brain |  |
| --- | --- | --- |
|  | WT | CD |
| **AEA** | 100 ± 1.73 | 101.54 ± 5.8 |
| **DEA** | 100 ± 6.6 | 97.60 ± 3.0 |
| **DHEA** | 100 ± 4.5 | 93.95 ± 4.9 |
| **2-AG** | 100 ± 3.5 | 111.1 ± 5.9 |
| **2-LG** | 100 ± 3.7 | 109.23 ± 6.1 |
| **2-OG** | 100 ± 2.9 | 100.05 ± 5.1 |

Next, we assessed cannabinoid receptor-mediated Gi/o protein activity by [^35^S]GTPγS autoradiography. CD mice exhibited a higher WIN55,212-2-stimulated [^35^S]GTPγS binding in several brain regions compared to WT mice, while basal activity was similar in both genotypes. The regions in CD mice with high G-protein coupling included the basolateral, lateral and central amygdala, piriform and visual cortex, olfactory bulb, and CA1 stratum oriens and CA1 stratum radiatum (Fig. 2c and d) (Supplementary Table 2). This increase was blocked in the presence of the CB1R antagonist rimonabant, but not with the CB2 antagonist SR144528 (Supplementary Fig. 2b). These results indicated that there was an increase in the functional coupling of CB1R to Gi/o proteins in several brain regions of CD mice. Interestingly, both CB1R density and CB1R- mediated Gi/o protein activity increased in different subregions of amygdala.

### JZL184 administration corrects behavioral impairment in CD mice

Previous reports have demonstrated that sub-chronic administration of JZL184, an irreversible inhibitor of the MAGL, promotes downregulation and G-protein uncoupling of CB1R (22, 23). We found that administration of JZL184 (8 mg/kg, i.p.) for 10 days significantly decreased the time that CD mice spent exploring stranger 1, reaching levels comparable to those displayed by WT mice during the sociability phase (Fig. 3a, left), whereas no significant changes were observed after a single dose of the drug (Supplementary Fig. 3). During the preference for novelty phase, CD mice treated with JZL184 showed a preference for social novelty similar to WT animals (Fig. 3a, middle). Notably, administration of JZL184 did not alter the exploration time of WT mice or the exploration time during the habituation phase (Fig. 3a, right). Furthermore, no changes were observed in locomotor activity after JZL184 administration in WT or in CD mice (Supplementary Fig. 4) discarding an overall effect of treatment that could bias exploratory activity of mice.

**Figure 3.**
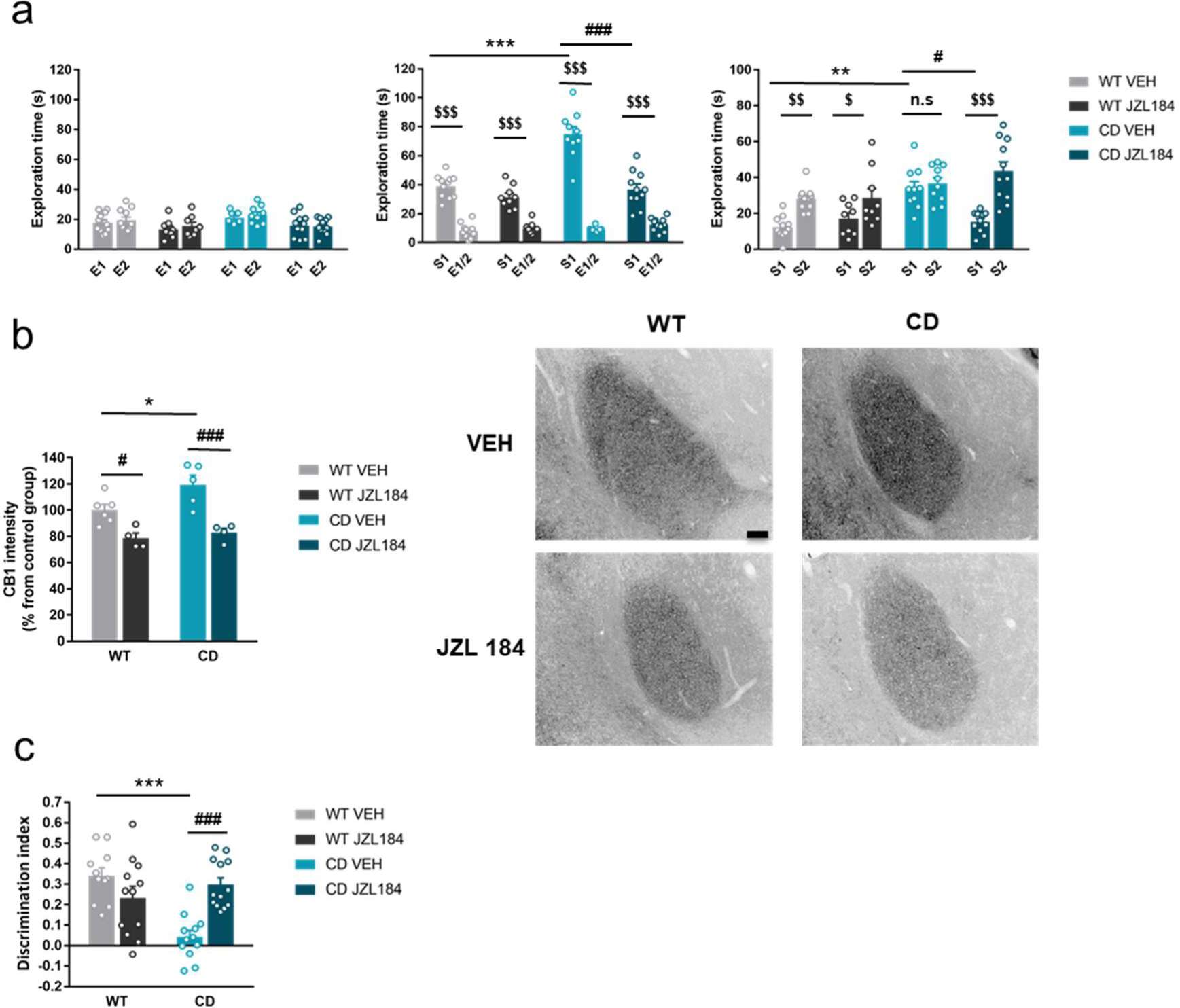
JZL184 treatment normalizes behavioral traits of CD mice. (a) Time spent exploring either empty compartments (E) or stranger mice (S) in the Vsocial-maze after 10 days of treatment of vehicle (VEH) or JZL184 (8 mg/kg) (WT VEH, n=11; WT JZL184, n=9; CD VEH, n=10; CD JZL184, n=11). Statistical significance was calculated by repeated measures ANOVA comparison. $ p<0.05; $$ p<0.01; $$$ p<0.001 (compartment effect); * p<0.05; *** p<0.001 (genotype effect); # p<0.05; ### p<0.001 (treatment effect). (b) Quantification and representative images of CB1R intensity in the basolateral amygdala of WT and CD mice after 10 days of treatment of vehicle (VEH) or JZL184 (8 mg/kg) (WT VEH, n=6; WT JZL184, n=4; CD VEH, n=5; CD JZL184, n=4). Scale bar=100 um. Statistical significance was calculated by calculated by Newman-Keuls post hoc test following two-way ANOVA. p*< 0.05 (genotype effect); # p< 0.05, ### p< 0.001 (treatment effect). (c) Discrimination index of WT and CD mice treated for 7 days with vehicle (VEH) or JZL184 (8 mg/kg) (WT VEH, n=11; WT JZL184, n=12; CD VEH, n=12; CD JZL184, n=13). Statistical significance was calculated by Newman-Keuls post hoc test following two-way ANOVA. p ***< 0.001 (genotype effect); ### p< 0.001 (treatment effect). Data are expressed as mean ± S.E.M

To determine whether sub-chronic administration of JZL184 at 8 mg/kg for 10 days induced changes in CB1R, we analyzed its density in amygdala. Immunofluorescence analysis showed that CD mice, as previously revealed by [^3^H]CP55,940 radioligand binding assay, presented an increased CB1R density in the basolateral amygdala. Moreover, we observed that sub-chronic JZL184 treatment decreased the density of CB1R in both WT and CD mice (Fig. 3b). Similar results were obtained by immunoblot using amygdala homogenates (Supplementary Fig. 5) confirming that 10 days of JZL184 treatment at 8 mg/kg downregulated CB1R levels in amygdala of CD mice. Given the role of the ECS in learning and memory processes (24), we studied the effect of JZL184 treatment over the recognition memory deficit of CD mice (Fig. 1d). Sub-chronic administration of JZL184 (8 mg/kg, i.p.) for 7 days (last administration 2 h before starting the training phase of the NORT), restored short-term memory impairment in CD mice (Fig. 3c). This improvement was not related to changes in the exploratory behavior, since total object exploration times did not change significantly among experimental groups (data not shown).

### JZL184 treatment has an impact on the cardiovascular phenotype of CD mice

We hypothesized that JZL184 treatment could impact the cardiovascular phenotype of CD mice. For this purpose, several anatomical parameters were evaluated. In agreement with previous descriptions of the CD model, mice presented an increase in the heart weight/body weight ratio (Fig. 4a) consistent with cardiac hypertrophy. In the morphometric analyses, wall-thickness of the left ventricle was significantly increased in CD mice compared to WT mice (Fig. 4b, left), which resulted in higher muscle proportion (Fig. 4b, right). Notably, administration of JZL184 for 10 days (8 mg/kg, i.p.) restored both the heart weight/body weight ratio and left ventricular wall-thickness (Fig. 4, a and b). Moreover, cardiac morphological changes showed an impact over cardiac function assessed by echocardiography. CD mice presented a slight reduction in left ventricular ejection fraction at baseline that was normalized after JZL184 (Fig. 4c), indicating that the treatment produced an improvement in cardiac function.

**Figure 4.**
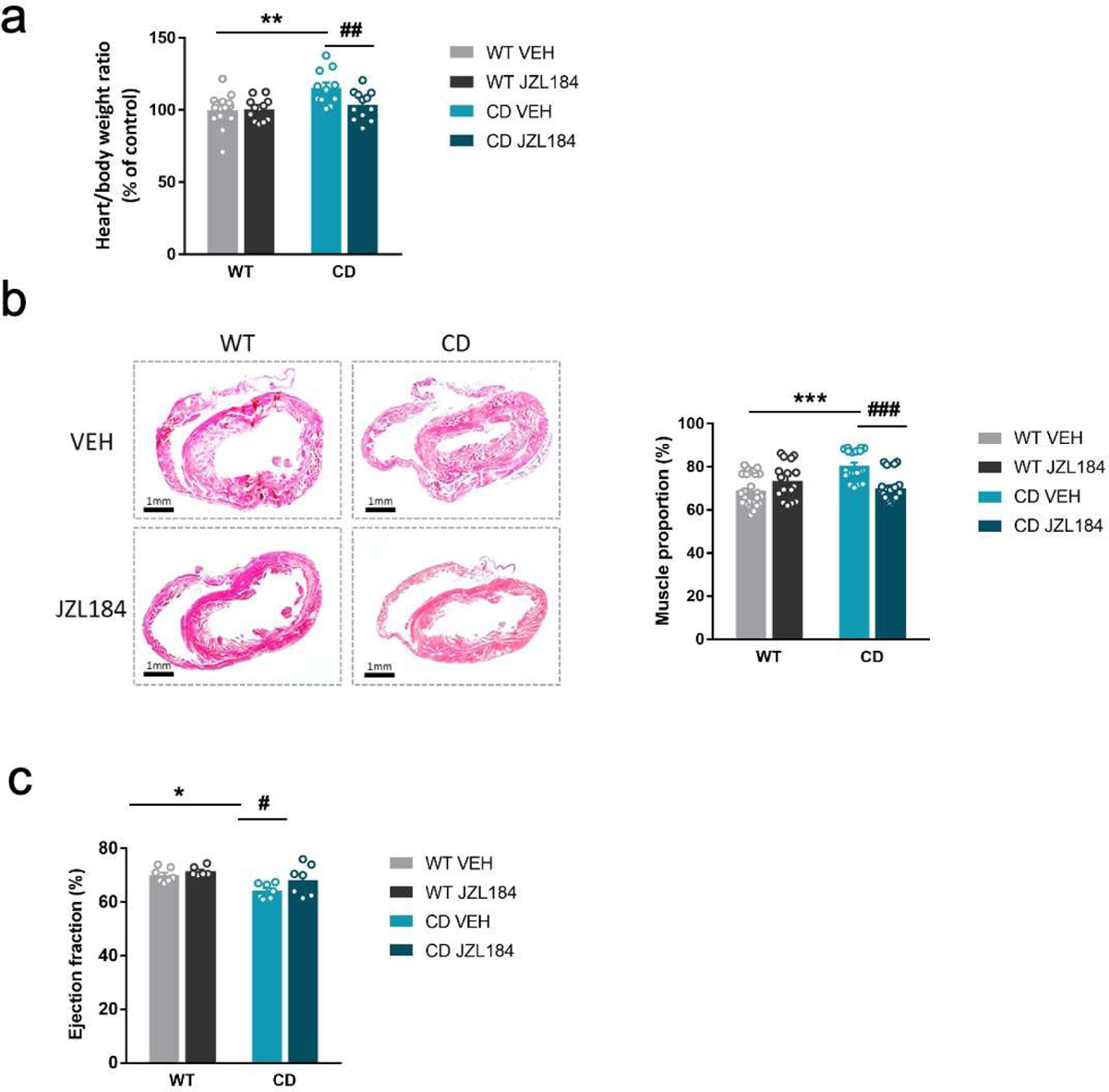
JZL184 administration regresses cardiac hypertrophy of CD mice. (a) Heart/body weight ratios obtained from WT and CD mice treated for 10 days with vehicle (VEH) or JZL184 (8 mg/kg) (WT VEH, n=14; WT JZL184, n=12; CD VEH, n=12; CD JZL184, n=13). (b) Representative haematoxylin-eosin stained transverse cardiac sections and quantification of muscle proportion from WT and CD mice treated for 10 days with vehicle (VEH) or JZL184 (8 mg/kg) (WT VEH, n=24; WT JZL184, n=16; CD VEH, n=20; CD JZL184, n=20 sections). (c) Ejection fraction (%) assessed by echocardiography from measurements performed on bidimensional images (WT VEH, n=8; WT JZL184, n=7; CD VEH, n=7; CD JZL184, n=7). Statistical significance was calculated by Newman- Keuls post hoc test following two-way ANOVA. * p<0.05; ** p<0.01; *** p<0.001 (genotype effect); # p<0.05; ## p<0.01; ### p<0.001 (treatment effect). Data are expressed as mean ± S.E.M.

### JZL184 treatment reverses cardiac transcriptional deficits of CD mice

Given the results on the cardiac phenotype in WBS, we performed a transcriptomic analysis on cardiac tissue to further explore the effects of JZL184 treatment in CD mice. For this purpose, we performed high-throughput RNA sequencing (RNA-seq) of heart samples from mice treated with vehicle or JZL184 for 10 days. Before analyzing differences in gene expression, a principal component analysis (PCA) was performed that revealed the samples used were informative with respect to the differences between experimental groups (Supplementary Fig. 6). After that, we first compared RNA-seq data between WT and CD treated with vehicle, and found 3,838 differentially expressed genes (DEGs) with a |log2 fold-change| > 0 and adjusted p values < 0.05 (Supplementary file 1) excluding the genes of the WBS critical region (Fig. 5a). Of these DEGs, 1,882 were upregulated and 1,956 downregulated, indicating a relevant alteration in cardiac gene expression in CD mice. Enrichment analysis identified Gene Ontology (GO) biological processes linked to the cardiovascular system including striated muscle cell development, muscle cell development, muscle system process, cardiac muscle contraction, heart contraction, heart process, contraction muscle cell development and muscle tissue progress among others (Fig. 5b). Then, we calculated the DEGs comparing vehicle-treated and JZL184-treated CD mice. This yielded 2,122 upregulated and 1,990 downregulated genes as a result of the treatment (Fig. 5c). Interestingly, overlap analysis revealed that 1,433 DEGs, 73 % of total downregulated in CD vehicle, were upregulated following JZL184 treatment (Fig. 5d). Enrichment analysis identified significant changes in several GO biological processes. Notably, among the 10 most significant GO biological processes, the majority were related to cardiovascular function including heart contraction, heart process, regulation of heart contraction, muscle system process, cardiac muscle contraction, muscle cell development and regulation of blood circulation (Fig. 5d). In addition, 1,262 DEGs, 67 % of total upregulated in CD vehicle, were downregulated after JZL184 treatment. Enrichment analysis identified much more diverse GO biological processes than those observed on the set of DEGs upregulated after JZL184 treatment, but also included some GO biological processes linked to the cardiovascular system such as regulation of vasculature development, endothelial cell migration and blood vessel endothelial cell migration (Fig. 5d). Moreover, when cardiovascular genes were only considered, the proportion of overlapping genes that reversed their expression as a result of treatment was higher than that described above: 79 % of downregulated and 69 % of upregulated in CD vehicle. Moreover, a concordance was observed in the direction and magnitude of the change in reverting cardiovascular genes (PearsonR=-0.9666, p-value=2.2e–16) (Figure 5e). Overall, these data indicated that JZL184 treatment may restore the normal expression of genes relevant for cardiovascular function that were either down- or upregulated in CD mice.

**Figure 5.**
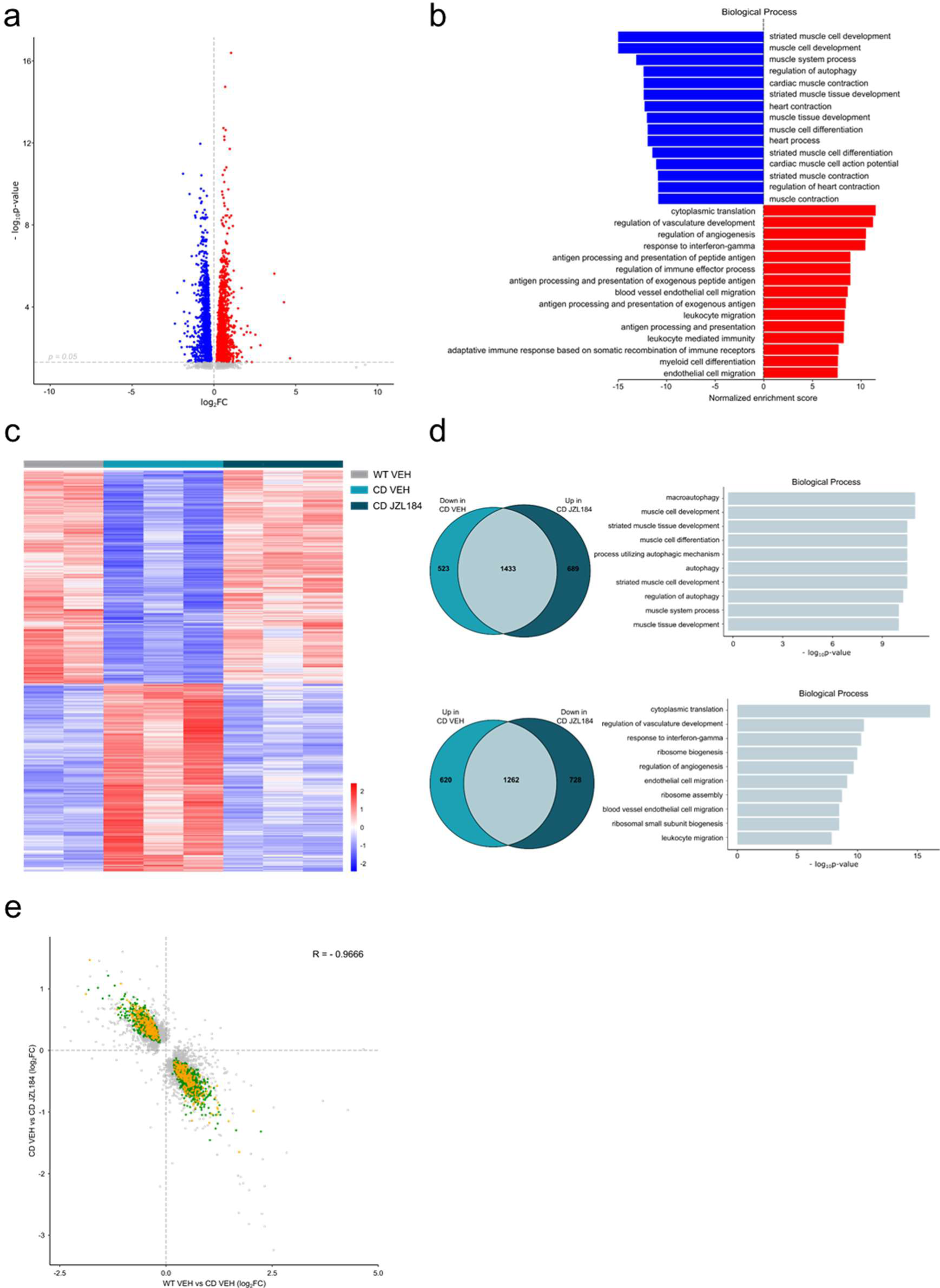
JZL184 treatment reversed alterations of the cardiac transcriptome observed in CD mice. (a) Volcano plot of differentially expressed genes (p<0.05 and |log2FC| > 0) between CD and WT mice. Red indicates relative increased expression and blue indicates relative decreased expression. (b) Gene ontology enrichment analysis for both up and down regulated genes in CD mice compared with WT. Most significant biological processes terms are represented for each group. (c) Heatmap showing the relative mRNA expression level of genes that reverted their expression in CD mice treated for 10 days with JZL184 (8 mg/kg) and CD or WT littermates treated with vehicle. (d) Venn diagrams and gene ontology enrichment analysis of genes that showed opposite differential expression in CD mice after 10 days treatment with JZL184 (8 mg/kg) compared with CD mice treated with vehicle. (e) Correlation plot of differentially expressed genes in CD mice treated for 10 days with JZL184 (8 mg/kg) and CD or WT littermates treated with vehicle. In green, genes with opposite differential expression between conditions, in orange reverted genes associated with cardiovascular function, in grey genes with no change. (WT VEH, n=2; CD VEH, n=3; CD JZL184, n=3).

## Discussion

The results in this study show that the ECS can be used as a target to improve characteristic behavioral and cardiac phenotypes in a relevant mouse model for WBS.

We first assessed social behavior in CD mice using an approach that allowed us to assess both sociability and preference for social novelty. In line with previous findings (13), we observed a significant increase in the sociability of these mice compared to WT mice. This phenotype resembles the human condition in which WBS subjects show higher social motivation (25, 26). In addition, we described for the first time that CD mice did not show preference for social novelty. This lack of social novelty preference is dependent on social stimuli since CD mice assessed on object novelty behaved similar to WT mice, and reflect a lack of habituation of CD mice to the previously encountered stranger mouse (13). This trait is reminiscent of the lack of habituation to faces observed through electrodermal measures in WBS individuals, which may cause that social stimuli appear continuously novel and interesting (27). These results improved the characterization of the social behavior of CD mice as a significant model of WBS.

The analysis of the main components of the ECS revealed an increased density and functional coupling to Gi/o proteins of CB1R in specific brain regions. Notably, density and functional coupling was increased in the amygdala, a region with major role in social behavior and where structural and functional abnormalities have been described in WBS patients (28–30).

Sub-chronic administration of the compound JZL184 at 8 mg/kg normalized the hypersociability and the lack of preference for social novelty of CD mice. The treatment did not have any effect over WT animals, revealing that the effects of JZL184 over social interactions were selective for the social phenotype of CD mice. Several studies have demonstrated that repeated administration of JZL184 at 8 mg/kg increase 2-AG levels by inhibiting MAGL in a long-lasting manner (22, 23, 31). The brain enhancement on 2-AG signaling resulting from JZL184 treatment for 10 consecutive days would be parsimonious with the decrease in CB1R density observed in the amygdala of CD mice. These results are in agreement with previous studies that have pointed to a downregulation of CB1R in chronic JZL184-exposed animals (22, 23, 32).

In the context of social behavior, previous results using JZL184 showed that a single systemic dose of this compound increased social play in juvenile rats (33) and social interaction in a mouse model of ASD, Shank3B–/– (15), and repeated social stress enhanced brain levels of 2-AG as well as decreased CB1R density (34). We did not observe a major effect over social behavior after a single dose of JZL184 in CD mice suggesting that changes in CB1R density and signaling after chronic exposure may be determinants of the effect observed over social behavior in CD mice. Together, we demonstrated that the sub-chronic administration of JZL184 at 8 mg/kg normalizes social abnormalities of CD mouse model. Atypical social functioning of WBS subjects predisposes to social vulnerability (35). In fact, WBS subjects have difficulties in peer interactions, maintaining friendships, and around 73 % have experienced social isolation (36). In addition, they have an increased risk to suffer psychiatric conditions that are not associated to the IQ range or to language disability (37) and seem related to the hypersocial phenotype (38, 39). Therefore, improvements in social functioning may have a beneficial effect over the quality of life of WBS subjects.

The treatment with the MAGL inhibitor JZL184 also restored short-term hippocampal-memory deficits of CD mice. In line with previous findings, our data also revealed that the JZL184 treatment did not have any effect over WT mice memory performance (31) and therefore, it is specific for the disorder context. Consequentially, our results on memory performance after JZL184 treatment could be further explored for cognitive restoration in the context of WBS. Notably, treatment with JZL184 may have additional benefits in WBS since it proved to have positive effects over the cardiac phenotype of CD mice. CD animals present a cardiovascular phenotype with cardiac hypertrophy present from an early age and accompanied by a decrease in the ejection fraction. We observed a significant normalization of the heart weight/body weight ratio and left ventricular wall-thickness after treatment with JZL184, indicating an improvement in heart hypertrophy. Moreover, ejection fraction was also improved after JZL184 treatment. This could be of major clinical relevance, given that the cardiovascular phenotype is the most life-threatening complication of the disorder, and constitutes a relevant new potential therapeutic approach for cardiovascular disorders.

CB1R are predominantly located in the cardiovascular centers of the brainstem and hypothalamus but also in myocardium, postganglionic autonomic nerve terminals and vascular endothelial and smooth muscle cells. Therefore, modulation of the ECS may have effects on the cardiovascular system by multiple mechanisms (40). Both agonists and antagonists of CB1R have shown to be beneficial for cardiovascular function in mouse models of different conditions (41–43) pointing that effects of the modulation of CB1R over cardiovascular systems seems to be highly dependent on the context (40). Our transcriptome analysis revealed that expression of the majority of genes that were down or up regulated in the CD mice, and were linked to cardiovascular function (including cardiac muscle contraction and development, and endothelial cell migration and vasculature development), were brought back to control levels after treatment with JZL184. On the other hand, we found an enrichment of genes related to macroautophagy/autophagy in CD samples after JZL184 treatment. These biological processes have been previously associated with improved cardiac function after heart failure (44). We also found a decrease in expression of genes involved in translation and ribosomal function, which could be associated with reduced hypertrophy (45). Further experiments may address the status of the different components of the ECS in the WBS context to better understand the mechanism of action of JZL184 over the cardiovascular system.

Taken together, the results of this study are of great importance given the few preclinical studies addressing potential treatments for WBS. In this regard, the modulation of the ECS may be an appropriate novel therapeutic strategy to tackle not only the social phenotype but also memory shortfalls and cardiovascular deficits in WBS.

## Methods

### Ethics

All animal procedures were conducted following ARRIVE (Animals in Research: Reporting In Vivo Experiments) guidelines (46) and standard ethical guidelines (European Union Directive 2010/63/EU) and approved by the local ethical committee (Comitè Ètic d’Experimentació Animal-Parc de Recerca Biomèdica de Barcelona, CEEA-PRBB).

### Animals

CD mice were obtained as previously described (13) and maintained on C57BL/6J background (backcrossed for nine generations). WT littermates were used as controls. Male mice aged between 8 and 16 weeks were used for experiments. In order to test social behavior juvenile (4 weeks old) male C57BL/6J mice were used as stranger mice.

Mice were housed in controlled environmental conditions (21 ± 1 °C temperature and 55 ± 10% humidity) and food and water were available ad libitum. All the experiments were performed during the light phase of a 12 h light/dark cycle (light on at 8 am; light off at 8 pm). Mice were habituated to the experimental room and handled for 1 week before starting the experiments. All behavioral experiments were conducted by an observer blind to the experimental conditions.

### Drug treatment

JZL184 (Abcam) was diluted in 15 % dimethyl sulfoxide (DMSO; Scharlau Chemie), 4.25 % polyethylene glycol 400 (Sigma-Aldrich), 4.25 % Tween-80 (Sigma-Aldrich) and 76.5 % saline. Rimonabant (Sanofi-Aventis) was diluted in 5 % ethanol, 5 % Cremophor-EL and 90 % saline. JZL184 and rimonabant were injected in a volume of 5 mL/kg and 10 mL/kg of body weight, respectively. Drugs were administered daily by i.p. injection 2 h prior behavioral testing.

### Behavioral tests

#### Sociability and preference for social novelty

Social behavior was performed in the V-social-maze (30-cm long × 4.5-cm wide × 15-cm height each corridor) as previously described (21). Briefly, the protocol consists of three phases: habituation (Phase I), sociability (Phase II) and preference for social novelty (Phase III). First, experimental mice were introduced into the central part of the V-maze where they freely explored the two empty chambers at the end of the corridors. This measurement is important to discard a possible bias for one particular chamber and it informs about the baseline activity of the mouse in the maze. Then, during the sociability phase an unfamiliar juvenile mouse assigned as stranger 1, was placed in one of the chambers (both sides were alternated during the experiments). The experimental mouse was allowed to explore both compartments for 5 min. The experimenter recorded the time that the experimental mouse spent exploring the empty chamber or the stranger 1. Finally, the preference for social novelty phase was performed just after the sociability session. A second novel juvenile mouse, assigned as stranger 2, was placed inside the previously empty chamber, while the stranger 1 remained inside the same chamber as in phase II. For 5 min, the experimental animal was allowed to explore the two strangers and the time spent exploring each stranger was recorded.

The procedure was performed in a sound-attenuated room with dim illumination 5-10 lux. A digital camera on top of the maze was used to record the sessions. Social exploration was considered when the experimental mouse directed the nose in close proximity (1 cm) to the vertical bars of the chambers. Mice that explored <10 s both mice were excluded from the analysis.

Acute treatment and last administration of the sub-chronic treatment (10 days) of JZL184 were performed 2 h before the V-social-maze test.

In order to assess the exploratory behavior towards objects in the same setting, the same procedure was performed using objects instead of juvenile stranger mice.

#### Locomotor activity

After 9 days of treatment with vehicle or JZL184, locomotor activity was assessed 2 h after the last administration. Spontaneous locomotor responses were assessed for 30 min by using individual locomotor activity boxes (9 cm width × 20 cm length × 11 cm high, Imetronic) in a low luminosity environment (5 lux). The total activity (number of horizontal movements) was detected by a line of photocells located 2 cm above the floor.

#### Novel object recognition test

The novel object recognition test (NORT) was performed as described before (47) in a V-shaped maze (V-maze). On day 1, mice were habituated to the empty V-maze for 9 min (habituation phase). On day 2, two identical objects (familiar objects) were presented at the end of each corridor of the V-maze and mice were left to explore for 9 min before they were returned to their home cage (familiarization phase). After 10 min, mice were placed back in the V-maze where one of the familiar objects was replaced by a new object (novel object) in order to assess memory performance (test phase). The time spent exploring each of the objects (familiar and novel) during the test session was computed to calculate a discrimination index (DI = (TIMEnovel-TIMEfamiliar)/(TIMEnovel+TIMEfamiliar)), defining exploration as the orientation of the nose toward the object within 2 cm from the object and with their nose facing it. A higher discrimination index is considered to reflect greater memory retention for the familiar object. Mice that explored <10 s both objects during the test session were excluded from the analysis. Drug administration was performed 2 h before the habituation and the training phases the 6^th^ and 7^th^ day of the sub-chronic treatment respectively.

### Endocannabinoid quantification by liquid chromatography–tandem mass spectrometry

The following N-acylethanolamines and 2-monoacylglycerols were quantified: N- arachidonoylethanolamine (AEA), N-docosatetraenoylethanolamine (DEA), N-docosahexa- enoylethanolamine (DHEA), 2-arachidonoylglycerol (2-AG), 2-linoleoylglycerol (2-LG) and 2- oleoylglycerol (2-OG). Its quantification was performed as described in (48). Briefly, half whole brains (231.4 ± 20.38 mg) (Mean ± S.D.) of mice were homogenized with 700 μL of 50 mM Tris-HCl buffer (pH 7.4): methanol (1:1) containing 25 μL of a mix of deuterated internal standards (5 ng/mL AEA-d4, 5 ng/mL DHEA-d4, 5 μg/mL 2-AG-d5, and 10 μg/mL 2-OG-d5) dissolved in acetonitrile. Afterwards, 5 mL of chloroform was added, and samples were shacked for 20 min and centrifuged at 1,700 g over 5 min at room temperature. Lower organic phase was evaporated under a stream of nitrogen, reconstituted in 100 μL of a mixture water:acetonitrile (10:90, v/v) with 0.1 % formic acid (v/v) and transferred to microvials for liquid chromatography analysis.

An Agilent 6410 triple quadrupole Liquid-Chromatograph equipped with a 1200 series binary pump, a column oven and a cooled autosampler (4 °C) were used to separate endocannabinoids. Chromatographic separation was carried out with a Waters C18-CSH column (3.1 x 100 mm, 1.8 μm particle size). The composition of the mobile phase was: A: 0.1 % (v/v) formic acid in water; B: 0.1 % (v/v) formic acid in acetonitrile. Gradient chromatography was used to separate endocannabinoids and related compounds and the ion source was operated in the positive electrospray mode. The selective reaction monitoring mode was used for the analysis. Quantification was done by isotope dilution with the response of the deuterated internal standards.

### Cannabinoid receptor autoradiography

Five fresh consecutive sections from brain of CD and WT mice (CD; n = 10 and WT; n=12) were dried and submerged in 50 mM Tris-HCl buffer containing 1 % of BSA (pH 7.4) for 30 min at room temperature, followed by incubation in the same buffer in the presence of the CB1R/CB2R radioligand, [^3^H]CP55,940 (3 nM) for 2 h at 37 °C. Nonspecific binding was measured by competition with non-labelled CP55,940 (10 µM) in another consecutive slice. The CB1R antagonist, SR141716A (1 µM) and the CB2R antagonist, SR144528 (1 µM), were used together with [^3^H]CP55,940 in two consecutive slices to check the CB1R or CB2R binding specificity. Then, sections were washed in ice-cold (4°C) 50 mM Tris-HCl buffer supplemented with 1 % BSA (pH 7.4) to stop the binding, followed by dipping in distilled ice-cold water and drying (4°C). Autoradiograms were generated by exposure of the tissues for 21 days at 4°C to β-radiation sensitive film together with [^3^H]-microscales used to calibrate the optical densities to fmol/mg tissue equivalent (fmol/mg t.e.).

### Labeling of activated Gi/o proteins by [^35^S]GTPγS binding assay

Brain samples from WT and CD mice were fresh frozen, cut into 20 µm sections, mounted onto gelatin-coated slides and stored (-25 °C) until used. Six fresh consecutive sections from CD (CD; n = 10) and wild type (WT; n = 12) mice were dried, followed by two consecutive incubations in HEPES-based buffer (50 mM HEPES, 100 mM NaCl, 3 mM MgCl2, 0.2 mM EGTA and 1 % BSA, pH 7.4) for 30 min at 30 °C. Briefly, sections were incubated for 2 h at 30 °C in the same buffer supplemented with 2 mM GDP, 1 mM DTT, and 0.04 nM [^35^S]GTPγS (PerkinElmer). The [^35^S]GTPγS basal binding was determined in two consecutive sections in the absence of agonist. The agonist-stimulated binding was determined in a consecutive brain section in the same reaction buffer in the presence of the CB1R/CB2R agonist, WIN55,212-2 (10 µM). The CB1R antagonist, SR141716A (1 µM) and the CB2R antagonist, SR144528 (1 µM), were used together with the agonist in two consecutive slices to check the specificity of the CB1R or CB2R functionality. Nonspecific binding was defined by competition with cold GTPγS (10 µM) in another section. Then, sections were washed twice in cold (4 °C) 50 mM HEPES buffer (pH 7.4), dried, and exposed to β-radiation sensitive film with a set of [^14^C] standards (American Radiolabelled Chemicals). After 48 h, the films were developed, scanned, and quantified by transforming optical densities into nCi/g tissue equivalence units using a calibration curve defined by the known values of the [^14^C] standards (ImageJ). Nonspecific binding values were subtracted from both agonist-stimulated and basal-stimulated conditions. The percentages of agonist-evoked stimulation were calculated from both the net basal and net agonist-stimulated [^35^S]GTPγS binding densities according to the following formula: ([^35^S]GTPγS agonist-stimulated binding x 100/[^35^S]GTPγS basal binding)-100.

### Tissue preparation for immunofluorescence

Immediately after social behavior assessment, a group of mice were deeply anesthetized by intraperitoneal injection of ketamine (100 mg/kg) and xylazine (20 mg/kg) mixture in a volume of 0.2 mL/10g of body weight. Subsequently, mice were intracardially perfused with 4 % paraformaldehyde in 0.1-M Na2HPO4/0.1-M NaH2PO4 buffer (PB), pH 7.5, delivered with a peristaltic pump at 19 mL/min flow for 3 min.

Afterwards, brains were extracted and post-fixed in the same solution for 24 h and transferred to a solution of 30 % sucrose in PB overnight at 4 °C. Coronal frozen sections (30 μm) of the basolateral amygdala (from Bregma: − 1.22 mm to − 1.82 mm) were obtained on a freezing microtome and stored in a solution of 5 % sucrose at 4 °C until used.

### Brain immunofluorescence and image analysis

Free-floating brain slices were rinsed in PB 0.1 M three times during 5 min with PB. Subsequently, brain slices were blocked in a solution containing 3 % donkey serum (DS) (D9663, Sigma-Aldrich) and 0.3 % Triton X-100 (T) in PB (DS-TPB) at room temperature for 2 h, and incubated overnight in the same solution with the primary antibody to CB1R (1:1,000, rabbit, Immunogenes) and neuronal nuclei (NeuN) (1:1,000, mouse, MAB377, Merck Millipore), at 4 °C. The next day, after 3 rinses in PB of 10 min each, sections were incubated at room temperature with the secondary antibody AlexaFluor-555 donkey anti-rabbit (for CB1R) (1:1,000, A-31572, Life Technologies, Thermo Fisher Scientific) and secondary antibody AlexaFluor-488 goat anti-mouse (for NeuN) (1:1,000, 115-545-003, Jackson ImmunoResearch) in DS-T-PB for 2 h. After incubation, sections were rinsed three times for 10 min each and mounted immediately after onto glass slides coated with gelatin in Fluoromont-G with 4′,6-diamidino-2-phenylindole (DAPI) (00-4959-52, Invitrogen, Thermo Fisher Scientific) as counterstaining.

Immunostained brain sections were analyzed with a ×10 objective using a Leica DMR microscope (Leica Microsystems) equipped with a digital camera Leica DFC 300FX (Leica Microsystems). The delimitation of basolateral amygdala area in each image was manually determined for quantification. The images were processed using the ImageJ analysis software. The mean intensity of the determined region was quantified using the automatic “measure” option of ImageJ. 2-4 representative images for each animal were quantified, and the average intensity of CB1R was calculated for each mouse. The data are expressed as a percentage of the control group (WT VEH). The displayed images were flipped for orientation consistency, adjusted for brightness and contrast and transformed to grey scale for display.

### Protein sample preparation

Immediately after social behavior assessment, amygdala and cardiac tissues were dissected from another group of mice. Tissues were frozen on dry ice and stored at − 80 °C un8l used, as previously reported. Samples from all animal groups, in each experiment, were processed in parallel to minimize inter-assay variations. The preparation of amygdala samples for total solubilized fraction, was performed as previously described (49). Frozen brain areas were dounce-homogenized in 30 volumes of lysis buffer (50 mM TrisHCl pH 7.4, 150 mM NaCl, 10 % glycerol, 1 mM EDTA, 10 μg/mL aprotinin, 1 μg/mL leupeptine, 1-μg/mL pepstatin, 1 mM phenylmethylsulfonyl fluoride, 1 mM sodium orthovanadate, 100 mM sodium fluoride, 5 mM sodium pyrophosphate, and 40 mM beta-glycerolphosphate) plus 1 % Triton X-100. After 10 min incubation at 4 °C, samples were centrifuged at 16,000 g for 20 min to remove insoluble debris. Protein contents in the supernatants were determined by DC-micro plate assay (Bio-Rad, Madrid, Spain), following manufacturer’s instructions.

### Immunoblot analysis

Anti-CB1R (1:500, rabbit, CB1-Rb-Af380-1, Frontier science) were detected using horseradish peroxidase-conjugated anti-rabbit antibody (1:15,000, Cell Signaling Technologies) and visualized by enhanced chemiluminescence detection (LuminataForte Western HRP substrate, Merck Millipore). Digital images were acquired on ChemiDoc XRS System (Bio-Rad) and quantified by The Quantity One software v4.6.3 (Bio-Rad). Optical density values for target proteins were normalized to Ponceau staining of the nitrocellulose membrane as loading control, and expressed as a percentage of control group (VEH treated mice).

### Echocardiogram

Echocardiograms were performed 2 h after the last administration of JZL184 sub-chronic treatment (10 d). Studies were carried out under general anaesthesia with isoflurane (2 %) using a Vivid IQ and a rodent-specific L8-18i-D Linear Array 5–15 MHz probe (General Electric Healthcare, Horten, Norway). Mice were placed in supine position on a continuously warmed platform to maintain body temperature, and all four limbs were fixed. Ultrasound gel was applied on the left hemithorax and hearts were imaged in parasternal short-axis projections. M-mode echocardiograms of the mid-ventricle were recorded at the level of the papillary muscles. The left ventricular end-diastolic (LVDD) and end-systolic (LVSD) internal diameters were measured in the M-mode recordings and were computed by the Teichholz formula into volumes as follows: LVDV=7*LVDD³/2.4 (where LVDV means LV end-diastolic volume) and LVSV=7*LVSD³/2.4+LVSD (where LVSV means LV end-systolic volume). LV ejection fraction (LVEF) was subsequently calculated as follows: LVEF= ((LVDV-LVSV)/LVDV)*100 and used as surrogate for LV systolic function as proposed by the American Society of Echocardiography (50). The average of 3 consecutive cardiac cycles was used for each measurement, with the operator being blinded to the group assignment.

### Heart histology and image analysis

Immediately after echocardiograms, mice were intracardiac perfused as described before. Afterwards, hearts were paraffin embedded and serial coronal sections (8 µm) were collected on a glass slide. Sections were stained with a regular hematoxylin-eosin protocol. Images of hematoxylin-eosin stained heart samples were obtained using visible light with an Olympus DP71 camera attached to an Olympus MVX10 MacroView Upright Microscope (zoom factor 1.25). The percentage of muscle was calculated by subtracting the lumen area from the total area occupied by the heart and indexing to the total area. Measures were performed with ImageJ software.

### RNA-seq analysis

Total RNA from heart was extracted following the standard protocol using TRIZOL reagent (Invitrogen) according to the manufacturer’s instructions. RNA-seq libraries were generated for Illumina sequencing with a HiSeq3000 instrument. The RNA-seq datasets were deposited on the Gene Expression Omnibus (GEO) and made publicly available through the accession GSE164257. Transcript abundances in Transcripts Per Million (TPM) were quantified with Salmon 0.7.2 using the Ensembl annotation GRCm38 v85. Read counts for genes were obtained from Salmon quantification with tximport (v 1.2.0) R package (51) and genes with less than 1 logCPM were filtered out. For the differential expression analysis, the count matrix was used with DESeq2 (v 1.26.0) Bioconductor package (52) as input directly from the tximport package. DESeq2 was used to estimate fold-changes and p-values for each gene between conditions and p-values were corrected using the Wald test procedure. The differentially expressed genes were selected based on a |log2 fold-change| > 0 and adjusted p-value < 0.05. Functional analysis was performed to examine the biological processes of the differentially expressed genes with clusterProfiler (v 3.14.3) Bioconductor package (53) Benjamini – Hochberg test was used to adjust the enrichment p-value for the gene ontology (GO) terms in each defined gene set using the expressed genes as background. Enriched GO terms were determined based on adjusted p-value < 0.05. Cardiovascular genes were selected from those genes with a described GO biological process term related with cardiovascular function.

Statistics Mice were randomly assigned to experimental groups. Sample size choice was based on previous studies (48, 54) and it is listed in figure legends for each experiment. Data were analyzed with Statistica Software using unpaired Student’s t-test and two-way ANOVA for multiple comparisons. Social interaction was analyzed by repeated-measures ANOVA with maze/genotype/treatment as between-subject factor and compartment as within-subject factor. Subsequent post hoc analysis (Newman-Keuls) was used when required (significant effect of factors or interaction between factors). Comparisons were considered statistically significant when p < 0.05. Outliers (± 2 S.D. from the mean) were excluded. The artwork was designed using GraphPad Prism 7. All results were expressed as mean ± S.E.M.

## Author contributions

A.N-R. participated in experimental design, conducted and analyzed behavioral and molecular experiments and wrote the manuscript. L.G-L. participated in experimental design, conducted and analyzed behavioral and molecular experiments and wrote the manuscript. P.O-R conducted and analyzed histological and molecular experiments. A.L-O. conducted and analyzed autoradiography experiments. L.dlR-R. analyzed and interpreted transcriptomic data and posted transcriptomic data in the public repository. A.M-S. conducted echocardiography analysis. M.R.-S. analyzed and interpreted transcriptomic data. A.P. performed endocannabinoid level measured. R.dlT. performed endocannabinoid level measures. R.M. participated in the supervision and experimental design. B.B-V. participated in electrocardiogram and histological interpretation. E.E. analyzed the transcriptomic data and provided input into their interpretation. R.R-P. analyzed autoradiography experiments. V.C. provided complete deletion (CD) mouse line, participated in histological analysis and discussed approaches. A.O. conceptualized, participated in experimental design, supervised and wrote the manuscript. All authors reviewed and approved the final version of the manuscript.

## Acknowledgments

We are grateful to Marta Linares, Dulce Real, Raquel Martín, Jolita Jancyte and Francisco Porrón for expert technical assistance and the Laboratori de Neurofarmacologia-NeuroPhar for helpful discussion.

## Conflict of interest statement

The authors declare that no conflict of interest exists.

---

**Supplementary Fig. 1.**
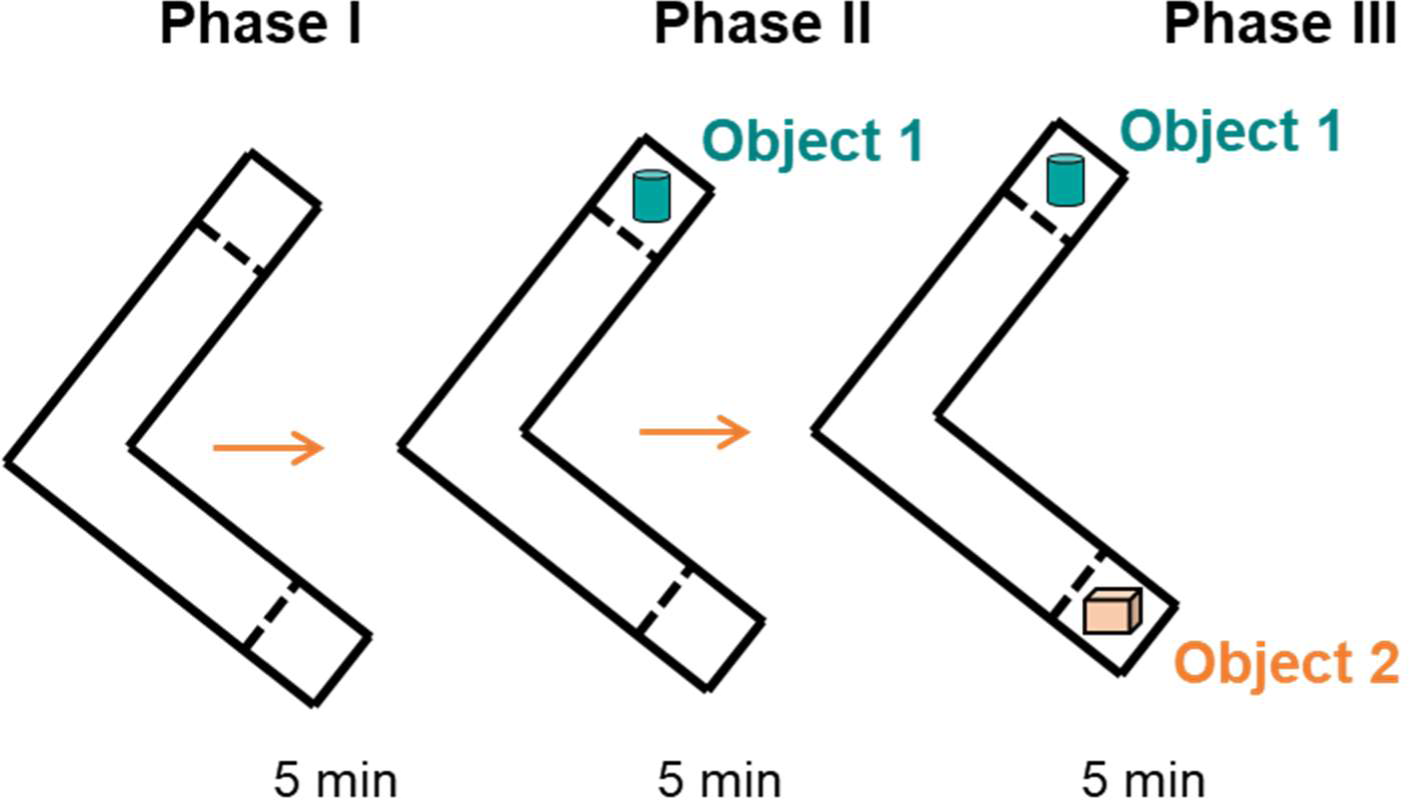
Schematic cartoon of the modification of the sociability and preference for social novelty procedure using unfamiliar objects instead of unfamiliar mice.

**Supplementary Fig. 2.**
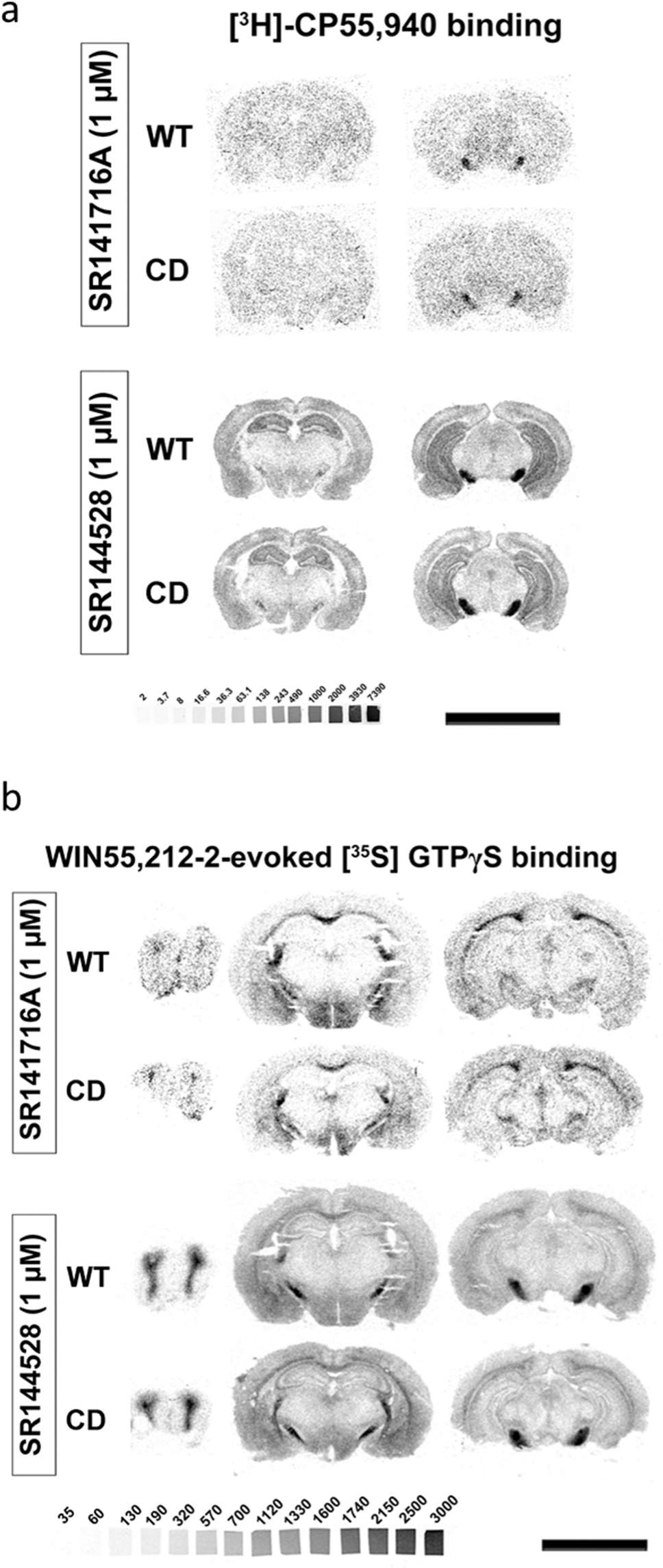
CD mice alterations in cannabinoid receptor density and activity are specific for CB1R. (a) Representative image of [^3^H]CP55,940 radioligand binding autoradiography. [^3^H]CP55,940 radioligand binding in brain slices was blocked with rimonabant but not with the CB2R antagonist SR144528. (b) Representative images of WIN55,212-2-evoked [^35^S]GTPγS binding. The increase in WIN55,212-2-stimulated [^35^S]GTPγS binding in brain slices was blocked in the presence of the CB1R antagonist rimonabant, but not with SR144528.

**Supplementary Fig. 3.**
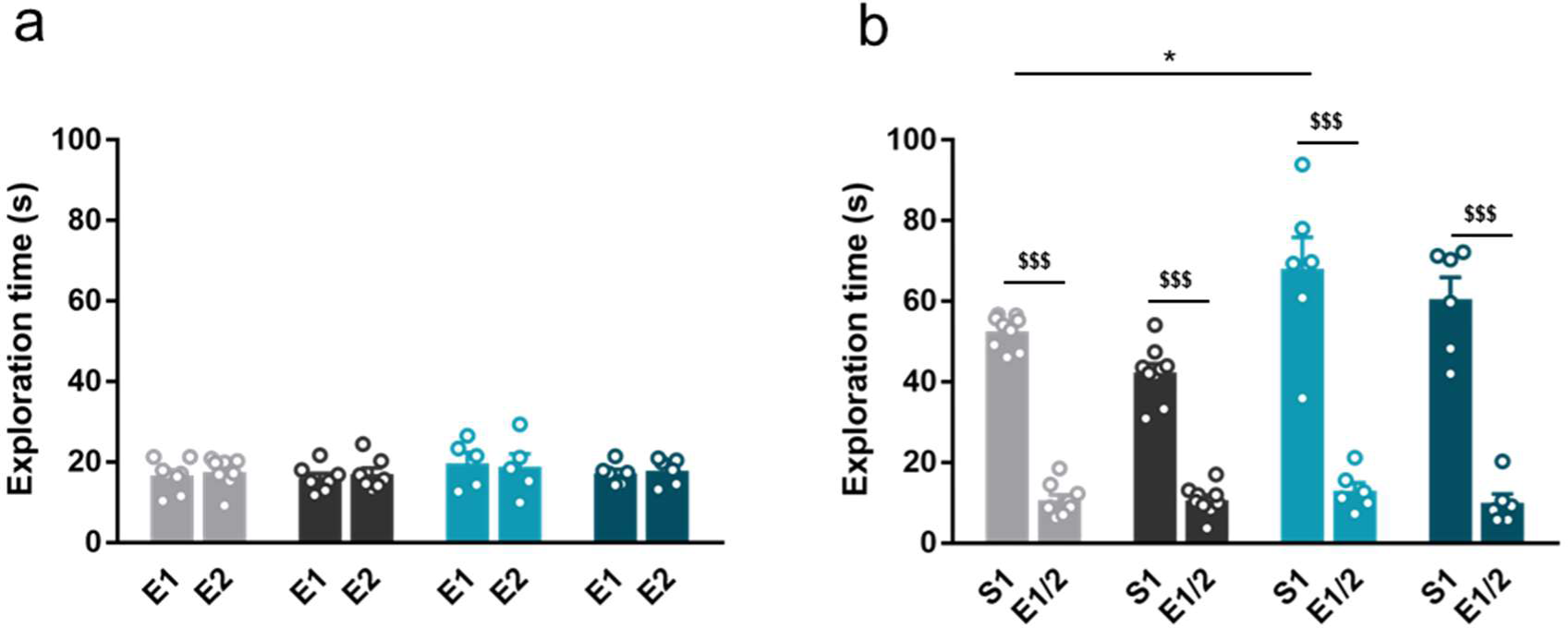
Acute administration of JZL184 does not normalise hypersociable phenotype in CD mice. (a) Time spent exploring both empty compartments (E). (b) Time spent exploring either empty compartments (E) or stranger mice (S) after one single of treatment of vehicle (VEH) or JZL184 (8 mg/kg) (WT VEH, n=8; WT JZL184, n=7; CD VEH, n=5; CD JZL184, n=6). Statistical significance was calculated by repeated measures ANOVA comparison. $$$ p<0.001 (compartment effect); * p<0.05 (genotype effect). Data are expressed as mean ± S.E.M.

**Supplementary Fig. 4.**
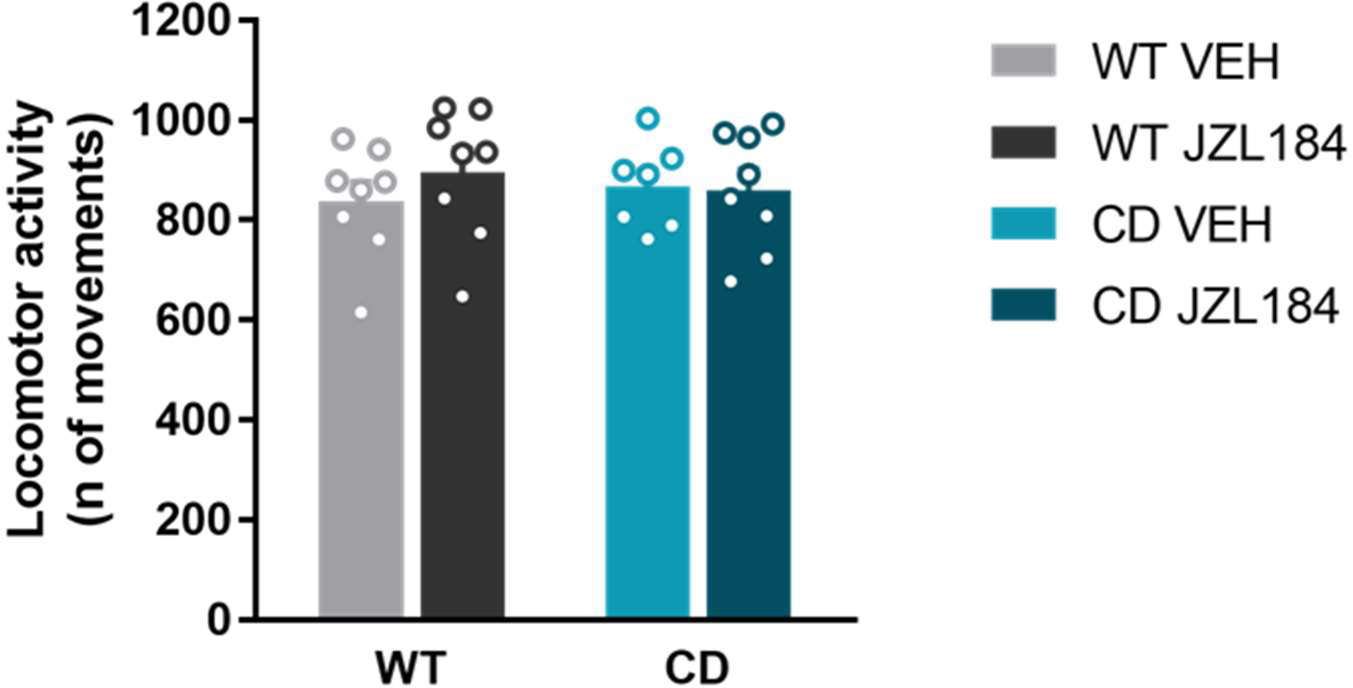
JZL184 treatment does not modify locomotor activity in WT and CD mice. Horizontal movements performed in locomotor activity boxes for 30 minutes by mice treated with vehicle (VEH) or JZL184 (8 mg/kg) (WT VEH, n=8; WT JZL184, n=8; CD VEH, n=8; CD JZL184, n=8). Statistical significance was calculated by two-way ANOVA. Data are expressed as mean ± S.E.M.

**Supplementary Fig. 5.**
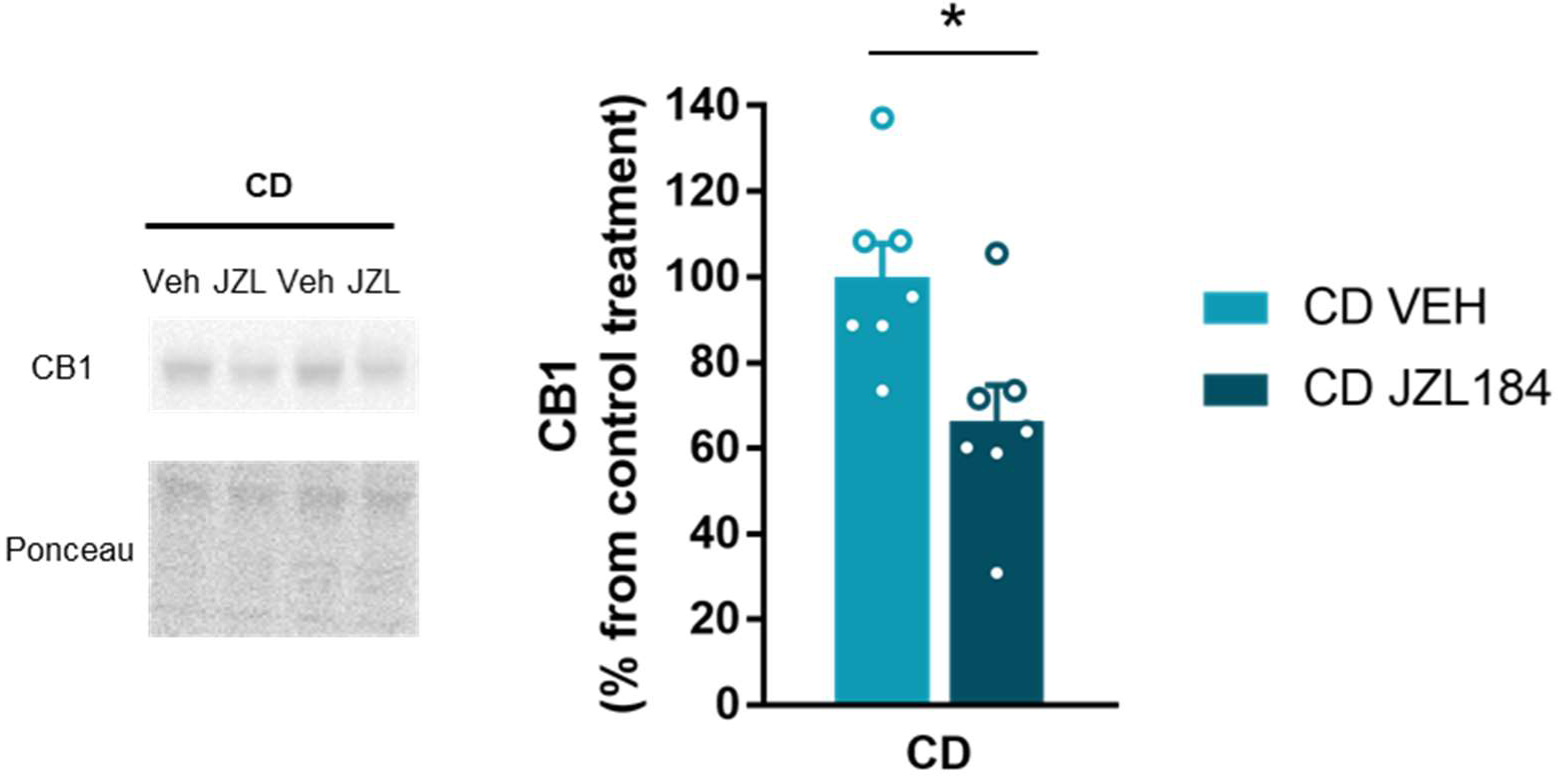
JZL184 treatment downregulates CB1R protein levels in the amygdala of CD mice (CD VEH, n=7; CD JZL184, n=7). Statistical significance was calculated by Student’s t- test. * p<0.05 (treatment effect). Data are expressed as mean ± S.E.M.

**Supplementary Fig. 6.**
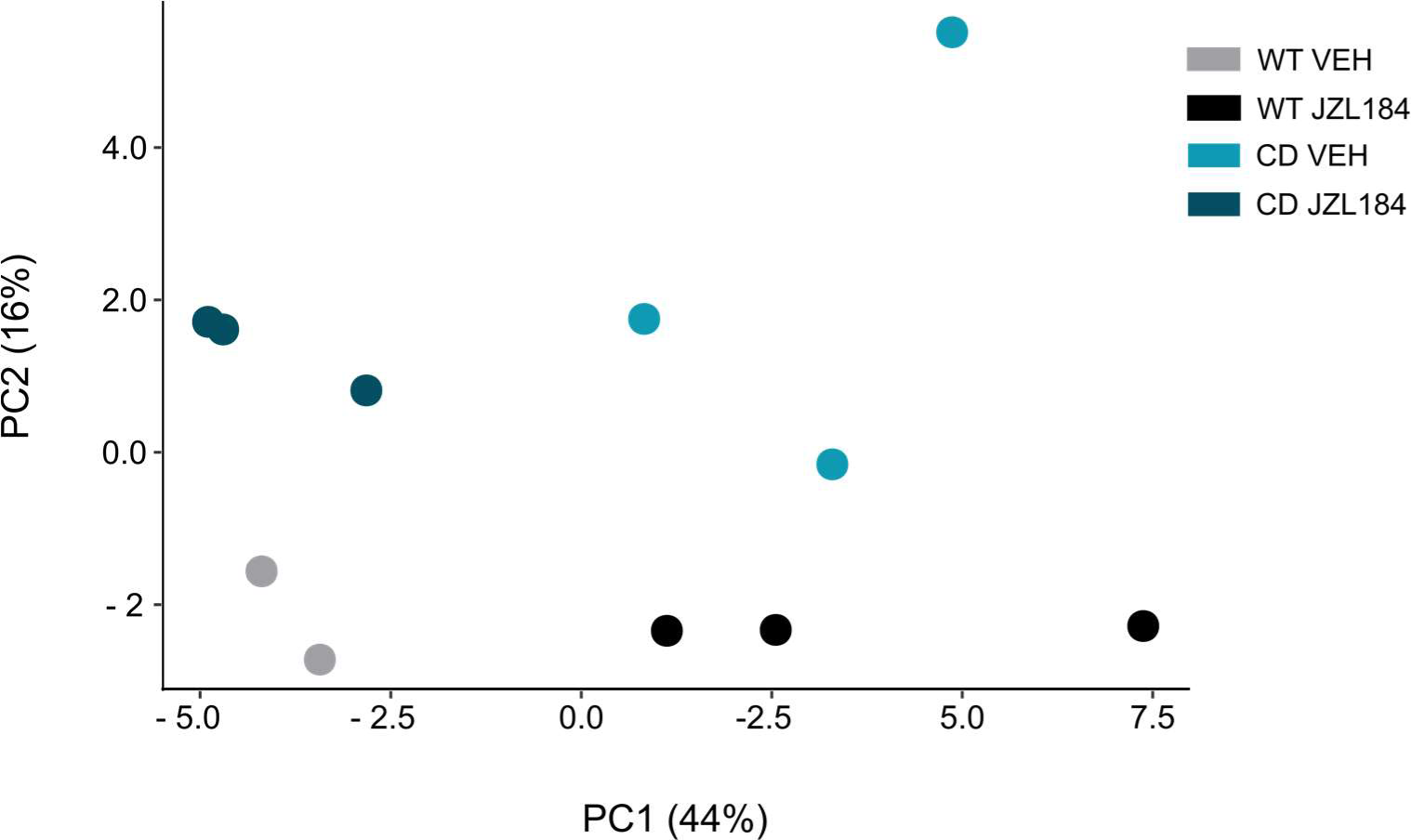
Principal component analysis (PCA) of sample-to-sample variation in gene expression.

**Supplementary Table 1.** [^3^H]CP55,940 binding of all analysed brain regions (WT, n=12; CD, n=10) in fmol/mg t.e. of CB1R. Statistical significance was calculated by Student’s t-test. * p<0.05; ** p<0.01; (genotype effect). Data are expressed as mean ± S.E.M.

|  | WT | CD |
| --- | --- | --- |
| ***Telencephalon*** |  |  |
| **Amygdala** |  |  |
| Anterior | 336 ± 21 | 368 ± 32 |
| Basolateral | 400 ± 21 | 494 ± 24** |
| Central | 464 ± 17 | 600 ± 54* |
| Lateral | 454 ± 26 | 496 ± 28 |
| Medial | 196 ± 18 | 233 ± 10 |
| Cortical amygdaloid nu | 552 ± 24 | 506 ± 43 |
| **Cortex** |  |  |
| Auditory | 243 ± 17 | 272 ± 13 |
| Cingular | 453 ± 36 | 479 ± 23 |
| Frontal | 578 ± 37 | 552 ± 36 |
| Ectorhinal | 414 ± 26 | 449 ± 32 |
| Entorhinal | 378 ± 20 | 419 ± 29 |
| Motor | 375 ± 29 | 411 ± 22 |
| Perirhinal | 384 ± 24 | 422 ± 24 |
| Piriform | 288 ± 13 | 334 ± 27 |
| Somatosensory | 298 ± 25 | 325 ± 21 |
| Visual | 298 ± 18 | 324 ± 22 |
| **Hippocampus** |  |  |
| CA1 |  |  |
| Oriens | 506 ± 31 | 531 ± 51 |
| Pyramidal | 443 ± 23 | 403 ± 38 |
| Radiatum | 674 ± 42 | 614 ± 54 |
| CA3 |  |  |
| Oriens | 743 ± 43 | 731 ± 59 |
| Pyramidal | 424 ± 37 | 350 ± 25 |
| Radiatum | 823 ± 47 | 796 ± 59 |
| Dentate Gyrus |  |  |
| Molecular | 610 ± 31 | 536 ± 50 |
| Polymorphic | 594 ± 28 | 470 ± 35* |
| Granular | 246 ± 25 | 176 ± 13* |
| Ventral subiculum | 1000 ± 42 | 986 ± 73 |
| **Basal ganglia** |  |  |
| Globus pallidus | 1958 ± 140 | 2061 ± 205 |
| Striatum | 577 ± 48 | 606 ± 44 |
| **Diencephalon** |  |  |
| Basal nucleus | 351 ± 25 | 426 ± 62 |
| Medial septum | 477 ± 23 | 488 ± 17 |
| **Rhinencephalon** |  |  |
| Olfactory bulb (glomerular) | 423 ± 17 | 460 ± 27 |
| **Rhomboencephalon** |  |  |
| Dorsal raphe | 310 ± 29 | 361 ± 30 |
| **Mesencephalon** |  |  |
| Periaqueductal Gray | 417 ± 39 | 477 ± 27 |
| Substantia nigra | 2248 ± 115 | 2350 ± 136 |
| **Metencephalon** |  |  |
| **Cerebellum** |  |  |
| Cerebelar gray matter | 1447 ± 80 | 1392 ± 92 |

**Supplementary Table 2.** [^35^S]GTPγS binding evoked by WIN55,212-2 (10 µM) of all analysed brain regions (WT, n=12; CD, n=10), expressed as percentage of stimulation over the basal binding. Statistical significance was calculated by Student’s t-test. * p<0.05; ** p<0.01; (genotype effect). Data are expressed as mean ± S.E.M.

|  | WT | CD |
| --- | --- | --- |
| *Telencephalon* |  |  |
| **Amygdala** |  |  |
| Anterior | 147 ± 18 | 163 ± 12 |
| Basolateral | 249 ± 25 | 349 ± 20 ** |
| Central | 95 ± 15 | 157 ± 24 * |
| Lateral | 202 ± 24 | 294 ± 34 * |
| Medial | 54 ± 8 | 87 ± 18 |
| Cortical amygdaloid nu | 188 ± 20 | 213 ± 19 |
| **Cortex** |  |  |
| Auditory | 112 ± 14 | 140 ± 12 |
| Cingular | 191 ± 17 | 224 ± 20 |
| Frontal | 216 ± 20 | 252 ± 30 |
| Ectorhinal | 219 ± 12 | 249 ± 22 |
| Entorhinal | 236 ± 17 | 281 ± 21 |
| Motor | 170 ± 12 | 190 ± 17 |
| Perirhinal | 232 ± 16 | 303 ± 31 |
| Piriform | 198 ± 19 | 257 ± 13 * |
| Somatosensory | 117 ± 10 | 141 ± 11 |
| Visual | 121 ± 13 | 173 ± 17 * |
| **Hippocampus** |  |  |
| CA1 |  |  |
| Oriens | 157 ± 13 | 226 ± 26 * |
| Pyramidal | 213 ± 18 | 263 ± 50 |
| Radiatum | 171 ± 12 | 232 ± 23* |
| CA3 |  |  |
| Oriens | 181 ± 23 | 199 ± 23 |
| Pyramidal | 240 ± 29 | 255 ± 25 |
| Radiatum | 237 ± 30 | 247 ± 38 |
| Dentate Gyrus |  |  |
| Molecular | 144 ± 23 | 127 ± 11 |
| Polymorphic | 224 ± 16 | 213 ± 14 |
| Granular | 243 ± 51 | 250 ± 31 |
| Ventral subiculum | 208 ± 20 | 204 ± 17 |
| **Basal ganglia** |  |  |
| Globus pallidus | 1533 ± 64 | 1627 ± 82 |
| Striatum | 203 ± 14 | 222 ± 16 |
| *Diencephalon* |  |  |
| Basal nucleus | 112 ± 9 | 147 ± 16 |
| Medial septum | 158 ± 11 | 161 ± 16 |
| *Rhinencephalon* |  |  |
| Olfactory bulb (glomerular) | 605 ± 34 | 757 ± 18 ** |
| *Rhomboencephalon* |  |  |
| Dorsal raphe | 64 ± 10 | 67 ± 7 |
| *Mesencephalon* |  |  |
| Periaqueductal Gray | 71 ± 9 | 101 ± 16 |
| Substantia nigra | 1760 ± 114 | 1694 ± 82 |
| *Metencephalon* |  |  |
| **Cerebellum** |  |  |
| Cerebelar gray matter | 380 ± 34 | 372 ± 37 |

## Notes

Funding A.N-R was the recipient of a predoctoral fellowship (Ministerio de Educación y Cultura, Spain), L.G-L. was supported by predoctoral fellowship by FPI (MINE- ICO/FEDER, EU). This study was supported by Ministerio de Economía, Innovación y Competitividad (MINECO), Spain (#RTI2018- 099282-B-I00B to A.O., #SAF2017-84060-R to R.M.; Generalitat de Catalunya, Spain (2017SGR- 669 to R.M.); Ministerio de Ciencia e Innovación (SAF2016-78508-R; AEI/MINEICO/FEDER, UE) to VC. Basque Government IT975-16 to the “Neurochemistry and Neurodegeneration” consolidated research group to R. R-P. ICREA (Institució Catalana de Recerca i Estudis Avançats, Spain) Academia to A.O. and R.M. Grant “Unidad de Excelencia María de Maeztu”, funded by the MINECO (#MDM-2014-0370); IPEP MdM 2017 to A.O. and E.E. FEDER, European Commission funding is also acknowledged.

### Competing Interest Statement

The authors have declared no competing interest.

